# Absolute and relative pitch processing in the human brain: Neural and behavioral evidence

**DOI:** 10.1101/526541

**Authors:** Simon Leipold, Christian Brauchli, Marielle Greber, Lutz Jäncke

## Abstract

Pitch is a primary perceptual dimension of sounds and is crucial in music and speech perception. When listening to melodies, most humans encode the relations between pitches into memory using an ability called relative pitch (RP). A small subpopulation, almost exclusively musicians, preferentially encode pitches using absolute pitch (AP): the ability to identify the pitch of a sound without an external reference. In this study, we recruited a large sample of musicians with AP (*AP musicians*) and without AP (*RP musicians*). The participants performed a pitch-processing task with a Listening and a Labeling condition during functional magnetic resonance imaging. General linear model analysis revealed that while labeling tones, AP musicians showed lower blood oxygenation level dependent (BOLD) signal in the inferior frontal gyrus and the presupplementary motor area — brain regions associated with working memory, language functions, and auditory imagery. At the same time, AP musicians labeled tones more accurately suggesting that AP might be an example of neural efficiency. In addition, using multivariate pattern analysis, we found that BOLD signal patterns in the inferior frontal gyrus and the presupplementary motor area differentiated between the groups. These clusters were similar, but not identical compared to the general linear model-based clusters. Therefore, information about AP and RP might be present on different spatial scales. While listening to tones, AP musicians showed increased BOLD signal in the right planum temporale which may reflect the matching of pitch information with internal templates and corroborates the importance of the planum temporale in AP processing. Taken together, AP and RP musicians show diverging frontal activations during Labeling and, more subtly, differences in right auditory activation during Listening. The results of this study do not support the previously reported importance of the dorsolateral prefrontal cortex in associating a pitch with its label.

## Introduction

Pitch is a primary perceptual dimension of sounds and plays a crucial role in music and speech perception (Plack et al. 2005). In humans, there exist differential mechanisms to encode pitches into memory. Most individuals encode pitches in relation to other pitches using an ability called relative pitch (RP). With the exception of individuals suffering from amusia (tone deafness), all humans are able to identify changes in pitch contour by making higher-lower judgements — even from a very young age (Plantinga and Trainor 2005). Trained musicians can also identify the exact musical interval (e.g., a perfect fifth) between pitches (McDermott and Oxenham 2008). A small subpopulation, almost exclusively comprised of musicians, preferentially encodes pitches in absolute terms (Miyazaki and Rakowski 2002). These musicians possess absolute pitch (AP), the ability to identify the pitch of a sound without an external reference (Zatorre 2003; Levitin and Rogers 2005; Deutsch 2013). In the following, musicians with AP are referred to as *AP musicians* and musicians without AP as *RP musicians*.

A cognitive theory of AP, the two-component model, postulates that AP consists of two separate processes: The first component (*pitch memory*) comprises long-term representations of pitches which presumably exist in all humans to some extent. The second component (*pitch labeling*) comprises the associations between the long-term pitch representations and meaningful labels (e.g., C#). These associations exist exclusively in AP musicians (Levitin 1994).

Although there has been a recent increase in neuroscientific AP research, the neural mechanisms underlying AP have been only partly identified. More than 20 years ago, it was first reported that AP musicians have a more pronounced left-right asymmetry of the planum temporale, a brain region located immediately posterior to Heschl’s gyrus on the superior temporal plane (Schlaug et al. 1995). Follow-up studies found that this asymmetry might be driven by a smaller size of the right planum temporale in AP musicians rather than by a larger left planum temporale (Keenan et al. 2001; Wilson et al. 2009; Wengenroth et al. 2014). With regard to the neurophysiology of AP, a seminal study used positron emission tomography (PET) to investigate pitch processing in AP and RP musicians (Zatorre et al. 1998). While listening to tones, AP musicians showed a unique increase in cerebral blood flow (CBF) in the left posterior dorsolateral prefrontal cortex (DLPFC). Because this region has been implicated in associative learning (Petrides et al. 1993), it was proposed that the CBF increase reflects the retrieval of the association between the pitch and its label from long-term memory. While labeling musical intervals, CBF increases in the posterior DLPFC were observed in both AP and RP musicians, but only RP musicians showed increases in the right inferior frontal gyrus (IFG). These increases were interpreted as reflecting working memory demands related to the RP ability (Zatorre et al. 1998).

In the general population, the prevalence of AP is roughly estimated to be less than one in 10,000 (Bachem 1955). Therefore, it is unsurprising that previous neuroscientific studies examining AP used small sample sizes. However, small samples result in low statistical power, which increases both the occurrence of false-negative and false-positive results (Button et al. 2013). As a consequence, previous neuroscientific AP studies reported inconsistent or even conflicting results. In this study, we aimed to counteract the statistical problems associated with small sample sizes by collecting and analyzing data from a large sample of musicians (n = 101). Using fMRI, we revisited the topic of pitch processing in AP and RP musicians. Similar to the aforementioned PET study, we employed a pitch-processing task comprising two experimental conditions (*Listening* vs. *Labeling*). Both AP and RP processing represented adequate strategies to solve the task due to its low difficulty (Itoh et al. 2005). Because individuals possessing AP preferentially encode pitches absolutely and non-possessors preferentially encode pitches relatively (Miyazaki and Rakowski 2002), the task allowed us to contrast AP and RP processing by comparing AP musicians with RP musicians.

According to the two-component model, AP musicians differ from RP musicians by having an association between the long-term representation of a pitch and its label (Levitin 1994). The retrieval of this pitch-label association might already occur during Listening and, to successfully perform the task, it must occur during Labeling (Zatorre et al. 1998). At the same time, AP musicians need not rely on working memory processes during Labeling (Itoh et al. 2005). For these reasons, we predicted smaller differences in AP musicians between Listening and Labeling both in BOLD signal responses and behavior. Because of their suggested role in AP processing, we expected an involvement of the posterior DLPFC and/or the planum temporale in AP musicians during Listening. Furthermore, we expected an involvement of the IFG in RP musicians during Labeling because of its association with working memory. Apart from conventional general linear model (GLM) analysis, we applied multivariate pattern analysis (MVPA) to the unsmoothed fMRI data to localize brain regions differentiating between AP and RP musicians. As a complement to GLM analysis, MVPA is sensitive to group-specific information being present in fine-grained voxel patterns which is not detectable using conventional analyses (Kriegeskorte and Bandettini 2007). Additionally, and independently from the other analyses, we investigated ROIs previously associated with AP for group differences which are homogeneous across a brain region but too subtle to be detected by voxel-wise analysis. ROI analysis provides more statistical power than the voxel-wise analyses due to the lower number of tests and thus, a less conservative correction for multiple comparisons (Poldrack 2007).

## Materials and Methods

### Participants

Fifty-two AP musicians and 50 RP musicians completed the pitch-processing task. Due to a technical error during the fMRI data export, one participant of the AP group was excluded, leaving the data of 101 participants for data analysis. The two groups were matched for sex, handedness, age, musical experience, and intelligence (see Table 1).

**Table 1.**
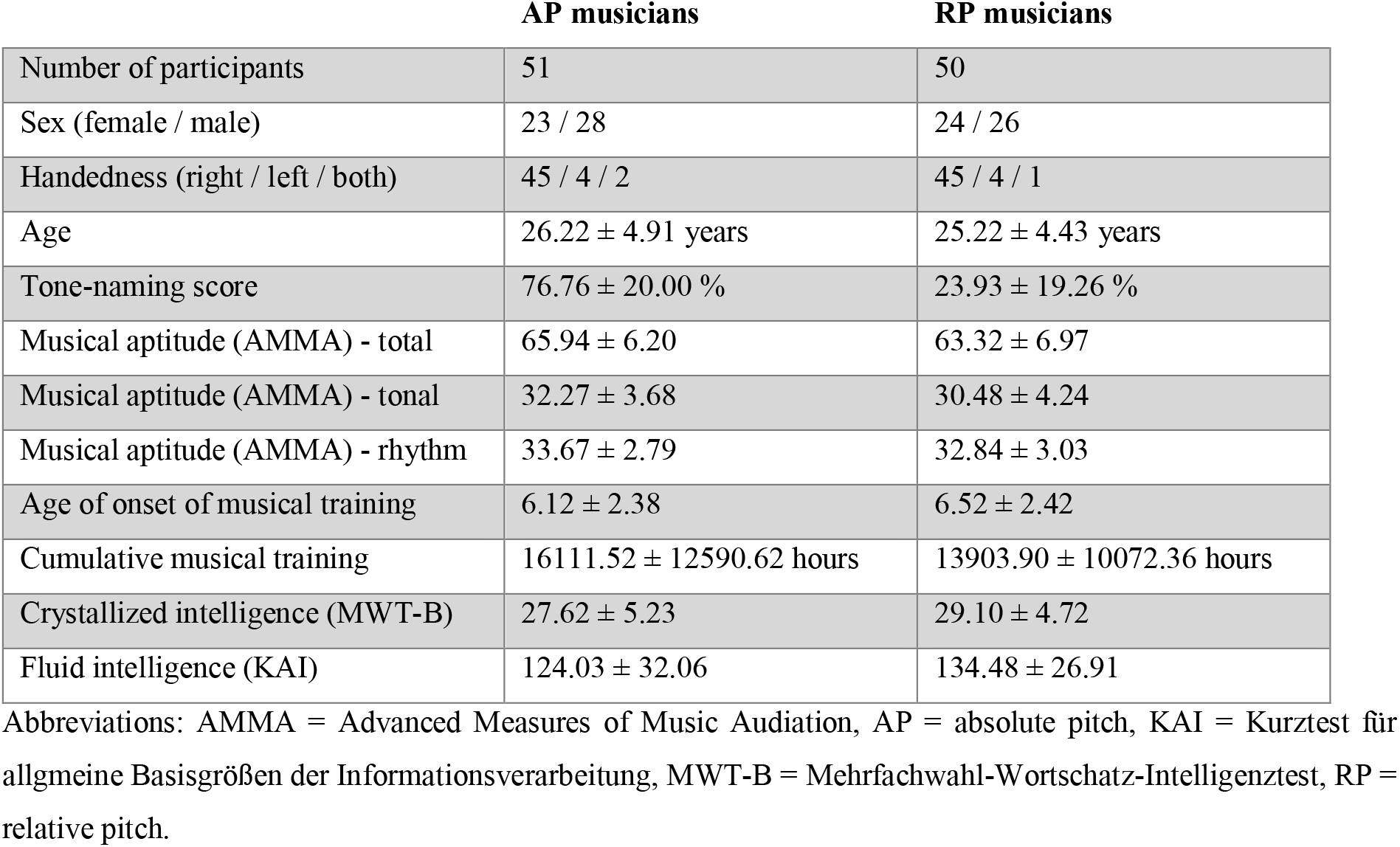
Participant characteristics. Continuous measures given as mean ± standard deviation.

|  | AP musicians | RP musicians |
| --- | --- | --- |
| Number of participants | 51 | 50 |
| Sex (female / male) | 23 / 28 | 24 / 26 |
| Handedness (right / left / both) | 45 / 4 / 2 | 45 / 4 / 1 |
| Age | 26.22 ± 4.91 years | 25.22 ± 4.43 years |
| Tone-naming score | 76.76 ± 20.00 % | 23.93 ± 19.26 % |
| Musical aptitude (AMMA) - total | 65.94 ± 6.20 | 63.32 ± 6.97 |
| Musical aptitude (AMMA) - tonal | 32.27 ± 3.68 | 30.48 ± 4.24 |
| Musical aptitude (AMMA) - rhythm | 33.67 ± 2.79 | 32.84 ± 3.03 |
| Age of onset of musical training | 6.12 ± 2.38 | 6.52 ± 2.42 |
| Cumulative musical training | 16111.52 ± 12590.62 hours | 13903.90 ± 10072.36 hours |
| Crystallized intelligence (MWT-B) | 27.62 ± 5.23 | 29.10 ± 4.72 |
| Fluid intelligence (KAI) | 124.03 ± 32.06 | 134.48 ± 26.91 |

Abbreviations: AMMA = Advanced Measures of Music Audiation, AP = absolute pitch, KAI = Kurztest für allgemeine Basisgrößen der Informationsverarbeitung, MWT-B = Mehrfachwahl-Wortschatz-Intelligenztest, RP = relative pitch.

Group assignment of the participants was based on self-report and confirmed by a tone-naming test (see below). Using both the information from self-report and a tone-naming test is advantageous because the assignment does not rely on an arbitrary cut-off concerning the tone-naming scores. In the rare case that a (potential) participant had indicated to be an AP musician in the initial online application form but then showed tone-naming scores around chance level (8.3%), we did not invite this participant for the imaging experiments in the laboratory. On the other hand, we did invite participants who had indicated to be RP musicians and then showed a high level of proficiency in tone-naming that was above chance level (and reiterated in the laboratory that they do not possess AP); please note that we did not regroup these participants as AP musicians. Furthermore, we statistically assessed if the group of RP musicians as a whole, and each RP musician individually, performed above chance level in the tone-naming test. On the group level, we found strong evidence that RP musicians performed better than chance (one sample t-test against 8.3%; *t*_(49)_ = 5.74, *P* < 10^−6^, Cohen’s *d* = 0.81). On the individual level, 56 % of the RP musicians performed above chance level according to a binomial test for each individual participant. Figure 1A shows the distribution of tone-naming scores. It is plausible that RP musicians performing above chance level used an internal reference (e.g., tuning standard 440 Hz) in combination with RP processing (or another yet unknown strategy) to solve the tone-naming test. Within RP musicians, tone naming did not correlate with age of onset of musical training (Pearson’s *r* = 0.06, *P* = 0.67) or with cumulative musical training (*r* = 0.17, *P* = 0.22).

**Figure 1.**
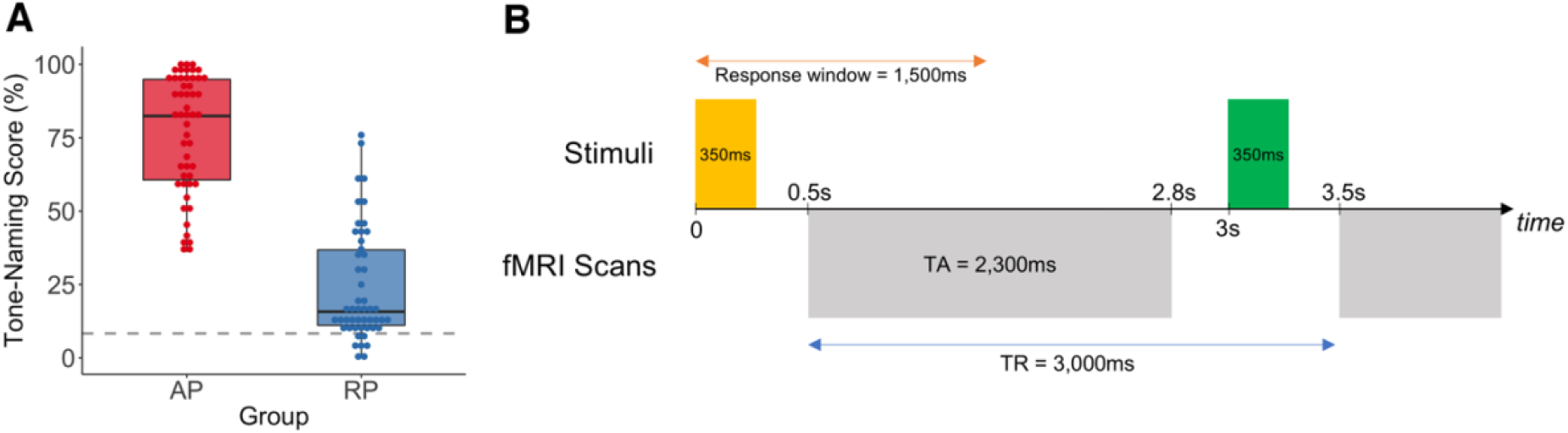
Tone-naming proficiency and fMRI task design. (*A*) Distribution of group-wise tone-naming scores. The dashed grey line represents chance level (8.3%). (*B*) A single trial of the fMRI task consisted of stimulus presentation, response, and scan acquisition. The TR was longer than the TA of a single scan, so stimuli could be presented in silence. AP = absolute pitch, RP = relative pitch, TA = acquisition time, TR = repetition time.

All participants were either music professionals, music students, or highly trained amateurs between 18 and 37 years. Participants were recruited in the context of a larger project investigating AP (Greber et al. 2018; Leipold et al. 2019; Brauchli et al. 2019; Burkhard et al. 2019), which involved multiple experiments using different imaging modalities (MRI, EEG). None of the participants reported any neurological, audiological, or severe psychiatric disorders, substance abuse, or other contraindications for MRI. The absence of hearing loss was confirmed by pure tone audiometry (ST20, MAICO Diagnostics GmbH, Berlin, Germany). Demographical data (sex, age, handedness) and part of the behavioral data (tone-naming proficiency, musical aptitude, and musical experience) was collected with an online survey tool (www.limesurvey.org). Self-reported handedness was confirmed using a German translation of the Annett questionnaire (Annett 1970). Musical aptitude was measured using the Advanced Measures of Music Audiation (AMMA) (Gordon 1989). Crystallized intelligence was estimated in the laboratory using the Mehrfachwahl-Wortschatz-Intelligenztest (MWT-B) (Lehrl 2005) and fluid intelligence was estimated using the Kurztest für allgmeine Basisgrößen der Informationsverarbeitung (KAI) (Lehrl et al. 1991). All participants provided written informed consent and were paid for their participation. The study was approved by the local ethics committee (www.kek.zh.ch) and conducted according to the principles defined in the Declaration of Helsinki.

### Sample Size Determination

We did not conduct a formal power analysis to determine the sample size for a given effect size and given power in advance of data acquisition. As data from AP musicians is extremely difficult to acquire due to their rarity (see Introduction), it was not possible to realistically plan for a specific number of participants to recruit. We rather recruited as many AP musicians as possible within a period of two years, given the limited financial and human resources available. The number of RP musicians was continuously updated to match the number of AP musicians already recruited at that time. With our final sample of about 50 participants per group, we had > 80% power to detect moderate to large effects (Cohen’s *d* > 0.6) in a two sample t-test setting. Please note that power analyses in the context of neuroimaging studies are difficult to perform (Mumford 2012), as effect sizes from previous smaller studies are probably inflated (Ioannidis 2008; Poldrack et al. 2017), and power analyses based on pilot studies are biased (Albers and Lakens 2018).

### Tone-Naming Test

Participants completed a tone-naming test to assess their tone-naming proficiency (Oechslin et al. 2010). The test was carried out online at home and participants were instructed to do the test in a silent environment where they could not be disturbed. During the test, 108 pure tones were presented in a pseudorandomized order. Each tone from C3 to B5 (twelve-tone equal temperament tuning, A4 = 440 Hz) was presented three times. The tones had a duration of 500 ms and were masked with Brownian noise (duration = 2000 ms), which was presented immediately before and after the tone. Participants were instructed to identify both the chroma and the octave of the tones (e.g., C4) within 15 s of tone presentation. To calculate a score of tone-naming proficiency, the percentage of correct chroma identifications was used. Octave errors were disregarded (Deutsch 2013). Therefore, the chance level identification performance was at 8.3%.

### Experimental Procedure

During fMRI scanning, participants performed a pitch-processing task (Zatorre et al. 1998; Itoh et al. 2005). The auditory stimuli used in the task consisted of three pure tones with different frequencies, and a segment of pink noise. The frequencies of the pure tones were 262 Hz (C4 in twelve-tone equal temperament tuning), 294 Hz (D4), and 330 Hz (E4). The pure tones and the noise segment had a duration of 350 ms with a 10 ms linear fade-in and a 50 ms linear fade-out. Therefore, all stimuli had an identical temporal envelope. The stimuli were created using Audacity (version 2.1.2, www.audacityteam.org). The pure tones and noise segments were presented via MRI-compatible headphones (NordicNeuroLab AS, Bergen, Norway).

The fMRI task was constructed as a rapid event-related design: Stimuli were presented in a randomized order and empty trials (without an auditory stimulus) were used to increase the efficiency of the design (Henson 2007). Within a trial, first, a stimulus (pure tone or noise segment) was presented for 350 ms; the participants were given 1,500 ms from stimulus onset to respond. Then, 500 ms after stimulus onset, we acquired a functional scan for 2,300 ms. Finally, the trial ended with 200 ms silence before the next trial began. Due to the prolonged repetition time (TR) of 3,000 ms between two scans in comparison with the acquisition time (TA) of 2,300 ms, the stimuli were presented in the silent period (700 ms) between the acquisitions of two subsequent scans. Therefore, there was no interference of scanner noise on the perception of the stimuli (Eden et al. 1999; Shah et al. 2000). The inter-trial interval (between two auditory stimuli) was varied using a jitter consisting of multiples of the TR (1–4 TRs). A visualization of the fMRI task is given in Figure 1B.

There were four runs in total. In each run, 39 pure tones (13 per chroma) and 39 noise segments were presented. The order of the stimuli was kept constant across the runs. Therefore, the auditory stimulation was identical in all runs. During the whole task, a black fixation cross on a gray background was presented on a screen. Stimulus presentation was controlled by Presentation software (version 17.1, www.neurobs.com). All stimuli and the stimulus presentation scripts are available online on the Open Science Framework (https://osf.io/ybghd/).

The task consisted of two experimental conditions: a Listening condition and a Labeling condition. These conditions only differed in the instructions given to the participants. In the Listening condition, participants had to press one response pad button (right middle finger) when they heard a pure tone, and another button (right index finger) when they had heard a noise segment. In the Labeling condition, participants had to label the pure tones by pressing one of three corresponding buttons on the response pad (right middle, ring, and little finger in response to C4, D4, and E4, respectively) and another button (right index finger) when they had heard a noise segment. The participants were instructed not to verbally respond and to respond as quickly and as accurately as possible. The accuracy of the responses and the response time were recorded via the response pad (4 button curved right, Current Designs INC, Philadelphia, PA, USA). Both conditions lasted for two runs each. The Listening condition always preceded the Labeling condition to avoid spillover effects from the Labeling onto the Listening condition. If the order had been the other way around, AP musicians might have been tempted to still covertly label the tones in the Listening condition.

### Statistical Analysis of Behavioral Data

In-scanner behavioral measures (response accuracy and response time) were analyzed in R (version 3.3.2, www.r-project.org). Separately for each measure, we performed a mixed-design ANOVA with a within-subject factor Condition (Listening vs. Labeling) and a between-subject factor Group (AP vs. RP). Subsequently, the two measures were separately compared within each condition using Welch’s t-tests. Next, we calculated differences in both measures by subtracting the Listening from the Labeling condition for each subject. These differences were then compared between the groups again using Welch’s t-tests. Finally, the differences were correlated with the tone-naming scores using the Pearson correlation coefficient. The significance level was set to *P* < 0.05. Generalized eta-squared (η^2^_G_ was used as an effect size for effects within an ANOVA and Cohen’s *d* for t-tests.

### Imaging Data Acquisition and Preprocessing

Imaging data was acquired on a Philips Ingenia 3.0 T MRI system (Philips Medical Systems, Best, The Netherlands), equipped with a commercial 15-channel head coil. Whole-brain functional images were acquired in four runs using a T2*-weighted gradient echo (GRE) echo planar imaging (EPI) sequence (scan duration of one run = 380 s). The T2*-weighted sequence had the following parameters: TR = 3000 ms, TA = 2300 ms, echo time (TE) = 35 ms, flip angle α = 90°, number of axial slices = 38, slice gap = 0.6 mm, slice scan order = interleaved, field of view (FOV) = 220 x 220 x 136 mm^3^, acquisition voxel size = 3.0 x 3.0 x 3.0 mm^3^, reconstructed voxel size = 2.75 x 2.75 x 3.6 mm^3^, reconstruction matrix = 80 x 80, number of dummy scans = 3, total number of scans = 122.

In addition, a whole-brain structural image was acquired using a T1-weighted GRE turbo field echo sequence (scan duration = 350 s). The T1-weighted sequence had the following parameters: TR = 8100 ms, TE = 3.7 ms, flip angle α = 8°, number of sagittal slices = 160, FOV = 240 x 240 x 160 mm^3^, acquisition voxel size = 1.0 x 1.0 x 1.0 mm^3^, reconstructed voxel size = 0.94 x 0.94 x 1.0 mm^3^, reconstruction matrix = 256 x 256. The whole scanning session lasted around 50 minutes and also involved resting-state fMRI and DTI. The results of these imaging modalities are discussed in other publications.

The functional images and the structural images were preprocessed using SPM12 (version 6906, www.fil.ion.ucl.ac.uk/spm/software/spm12). The following preprocessing steps were performed in succession using default settings unless otherwise stated: (i) Slice time correction. (ii) Motion correction by a rigid body transformation using six parameters (three translations and three rotations). We did not use unwarping as we had not collected data to correct geometrical distortions caused by susceptibility-induced magnetic field inhomogeneities. (iii) Coregistration of the structural image to the mean functional image. (iv) Segmentation and bias field correction of the structural image and estimation of the deformation field to map the image to the T1-weighted MNI152 template. (v) Normalization of the functional images using the estimated deformation field. (vi) Interpolation to an isotropic voxel size of 3.0 mm. (vii) Smoothing of the functional images with an 8 mm full width at half maximum (FWHM) three-dimensional Gaussian kernel. The quality of the normalization was visually inspected to confirm proper execution.

### GLM Analysis

Subject-wise first-level analysis was performed in SPM12. The voxel-wise BOLD signal time series was modeled using a GLM. The first-level design matrix contained, for each run separately, two regressors of interest (onsets of pure tones, onsets of noise segments) and one regressor of no interest (onsets of button presses). These regressors were modeled by convolving delta functions with the canonical double-gamma hemodynamic response function (HRF). Furthermore, we included the six motion parameters estimated during preprocessing as nuisance regressors and applied a high-pass filter (cutoff = 128 s) to remove low-frequency drifts. The following first-level contrasts of interest were calculated: Tones _Listening_ > Noise _Listening_ and Tones _Labeling_ > Noise _Labeling_. Following the logic of cognitive subtraction, these contrasts reflect BOLD signal increases associated with pitch processing.

Second-level random effects analysis was performed using non-parametric permutation tests as implemented in SnPM13 (www.warwick.ac.uk/snpm). Permutation tests depend on fewer assumptions than standard parametric approaches and provide an exact control of the family-wise error (FWE) rate (Nichols and Holmes 2002; Eklund et al. 2016). For the second-level analysis, we used a 2 x 2 mixed factorial design to investigate the interaction between Group (AP vs. RP) and Condition (Listening vs. Labeling). To facilitate the interpretation of the interaction, difference images were created for each subject by subtracting the contrast image of the Listening condition (Tones _Listening_ > Noise _Listening_) from the contrast image of the Labeling condition (Tones _Labeling_ > Noise _Labeling_). These difference images were entered in SnPM13 as inputs for a two sample t-test to compare AP and RP musicians (cluster-wise inference, 10000 permutations, cluster defining threshold (CDT) *P* < 0.001). An anatomically defined mask was used to restrict the search space of the analysis to a priori defined brain regions previously associated with AP and RP processing. To create this mask, we used probability maps of the following bilateral brain regions included in the Harvard-Oxford cortical atlas (http://fsl.fmrib.ox.ac.uk/fsl/fslwiki/Atlases). (i) Heschl’s gyrus, (ii) planum temporale, (iii) planum polare, (iv) superior temporal gyrus (anterior and posterior division), (v) superior frontal gyrus, (vi) middle frontal gyrus, (vi) inferior frontal gyrus (pars opercularis and pars triangularis), (vii) superior parietal lobule, (ix) gyrus supramarginalis (anterior and posterior division), and (x) angular gyrus. The probability maps were then combined, thresholded and binarized at 10% probability using the utility fslmaths (http://fsl.fmrib.ox.ac.uk/fsl/fslwiki/Fslutils). Using a mask to restrict the search space alleviates the problem of multiple comparisons as less voxels are tested for an effect (Poldrack 2007). This particular mask furthermore reflects prior knowledge that has been accumulated about AP and RP in many studies over the years. Structural and functional alterations of auditory regions in the superior temporal cortex have been repeatedly linked to AP processing (Schlaug et al. 1995; Keenan et al. 2001; Wilson et al. 2009; Jäncke et al. 2012; Schulze et al. 2013; Wengenroth et al. 2014; Kim and Knösche 2016, 2017; McKetton et al. 2019; Brauchli et al. 2019). Dorsal and ventral frontal areas have been associated with both AP and RP (Zatorre et al. 1998; Ohnishi et al. 2001; Bermudez et al. 2009; Wengenroth et al. 2014; Dohn et al. 2015; Brauchli et al. 2019), and there is evidence that parietal areas contribute to AP (Loui et al. 2012; Brauchli et al. 2019) and RP (Schulze et al. 2009; Foster and Zatorre 2010b, a).

Two follow-up analyses with the same mask were performed. To determine the effects of condition within each group, we entered the difference images as inputs for a one sample t-test for each group separately (cluster-wise inference, 10000 permutations, CDT *P* < 0.001). To determine the effects of group within each condition, we entered the first-level contrast images (Tones _Listening_ > Noise _Listening_, Tones _Labeling_ > Noise _Labeling_) as inputs for a one sample t-test for each condition separately (cluster-wise inference, 10000 permutations, CDT *P* < 0.001). The significance level for all analyses was set to α = 0.05, FWE-corrected for multiple comparisons.

Additionally, we performed a GLM-based whole-brain analysis to explore effects located outside of brain regions previously associated with AP or RP. This exploratory analysis extended the search space to all brain regions of the Harvard-Oxford cortical and subcortical atlases (excluding the cerebral white matter, the brain stem, and the lateral ventricles). In this whole-brain analysis, we employed the same second-level analysis steps as described above for the restricted analysis.

### MVPA

We carried out a specific type of MVPA, namely searchlight analysis as implemented in PyMVPA (version 2.6.1, www.pymvpa.org) to detect brain regions containing fine-grained BOLD signal patterns which differentiated between AP and RP musicians (Kriegeskorte et al. 2006; Etzel et al. 2013). Due to the high computational demands, all analyses were carried out on the ScienceCloud of the University of Zurich (www.s3it.uzh.ch). Searchlight analysis, sometimes called information-based brain mapping, builds a map of voxels which are informative regarding group status (searchlight analysis can also be used to analyze information about different stimuli or experimental conditions). A machine learning classifier uses local BOLD signal patterns to classify the participants as belonging to one of the two groups. Brain regions which contain clusters of informative voxels are differentially activated in the two groups (Kriegeskorte et al. 2006; Kriegeskorte and Bandettini 2007).

Searchlight analysis was performed on the unsmoothed functional images. To some extent, smoothing removes the fine-grained patterns of activation which were here analyzed for information about group status (Kriegeskorte and Bandettini 2007). Analogous to the GLM analysis, two first-level contrasts were computed in SPM12 (this time using the unsmoothed images): Tones _Listening_ > Noise _Listening_ and Tones _Labeling_ > Noise _Labeling_. In addition, we again calculated a difference image for each subject by subtracting the contrast image of the Listening condition from the contrast image of the Labeling condition.

In total, we performed three searchlight analyses using the different images (difference images, Listening contrast images, Labeling contrast images) as inputs. In all analyses, a sphere was moved across all voxels of the anatomically defined mask that was also used in the GLM analysis. Each sphere had a radius of three voxels (9 mm) and consisted of one center voxel and (at most) 122 surrounding voxels. In every sphere, a linear support vector machine (C = 1) was trained and tested using a 5-fold cross-validation. For the cross-validation, the input images were pseudorandomly partitioned into five chunks under the restriction that each chunk contained the same number of images of AP musicians and RP musicians. One chunk contained 11 images of AP musicians (instead of 10), because our analyzed sample included 51 AP and 50 RP musicians. The average classification accuracy of the five folds was written in the location of the center voxel to create a map of classification accuracies (i.e. an information map).

To assess the statistical significance of informative clusters, we used non-parametric permutation testing (Nichols and Holmes 2002). For this purpose, each of the three searchlight analyses was repeated with permuted group labels (10000 permutations). For every iteration, the group labels were randomly permuted within each chunk. We used this restriction to balance the number of images per group in each chunk. The resulting permutation set was fixed for the whole searchlight analysis (i.e. across all center voxels of the mask) to preserve the spatial dependency between neighboring center voxels (Stelzer et al. 2013). All properties of the searchlight analyses with the permutated labels were identical to the analyses with the real labels (e.g., classifier parameters, cross-validation scheme). The permutation procedure resulted in a null distribution of 10000 information maps.

Next, both the empirical information map (created with the real labels) and the null information maps (created with the permuted labels) were thresholded with a CDT of *P* < 0.001 using custom MATLAB R2016a functions. Subsequently, we formed clusters of the above-threshold voxels using CoSMoMVPA (version 1.1.0, www.cosmomvpa.org). The maximum cluster size of each null information map was extracted to form a null distribution of cluster sizes. Finally, the *P* value of the clusters in the empirical information map was calculated as the proportion of cluster sizes under the null distribution that were larger than the empirical cluster size. The significance level was set to α = 0.05, FWE-corrected.

### ROI Analysis

In addition to the voxel-wise GLM and searchlight analyses, the mean BOLD signal changes in a priori defined ROIs were compared between groups using MarsBaR (version 0.44, www.marsbar.sourceforge.net). We defined four ROIs which have been previously associated with AP processing: left planum temporale (Schlaug et al. 1995; Wilson et al. 2009), right planum temporale (Keenan et al. 2001; Wilson et al. 2009; Wengenroth et al. 2014), left DLPFC (Zatorre et al. 1998; Ohnishi et al. 2001; Bermudez and Zatorre 2005), and right DLPFC (Bermudez and Zatorre 2005).

The ROIs were created as spheres (radius = 10 mm) based on MNI coordinates. We used anatomically defined coordinates for the planum temporale and functionally defined coordinates for the DLPFC, because the planum temporale can be delineated by anatomical landmarks, whereas the DLPFC is primarily a functional region. The coordinates of the left (x = −44, y = −34, z = 11) and right planum temporale (x = 41, y = −31, z = 15) were derived from the Harvard-Oxford cortical atlas planum temporale probability map by choosing the voxel with the highest probability in the left and the right hemisphere. The coordinates of the left DLPFC (x = −40, y = 9, z = 42) were taken from a seminal study investigating pitch processing in AP, which was the first to associate this brain region with the retrieval of the pitch-label association while AP musicians were listening to tones (Zatorre et al. 1998). The original study reported the coordinates in Talairach space, so we transformed the coordinates into MNI space (Lacadie et al. 2008). The coordinates of the left hemispheric region were flipped at the midsagittal plane to derive the coordinates of the right DLPFC (x = 40, y = 9, z = 42). For each subject and ROI, we extracted first-level contrast values from the Listening condition (Tones Listening > Noise Listening). For each ROI, these contrast values were compared between AP and RP musicians using Welch’s t-tests in R. The significance level was set to α = 0.0125, FWE-corrected for multiple ROIs.

At the request of a reviewer, we conducted a supplemental ROI analysis for the bilateral Heschl’s gyrus. Recent studies have implicated both structure and function of Heschl’s gyrus in AP processing (Wengenroth et al. 2014; McKetton et al. 2019; Brauchli et al. 2019). Coordinates for left (x = −44, y = −24, z = 11) and right Heschl’s gyrus (x = 42, y = −20, z = 9) were anatomically defined, analogous to the planum temporale, by choosing the voxel with the highest probability per hemisphere of the Harvard-Oxford cortical atlas Heschl’s gyrus probability map. In line with the exploratory character of this analysis, here, we used a significance level of α = 0.05, uncorrected.

## Results

### Behavior

Demographical and behavioral characteristics of the AP musicians (n = 51) and the RP musicians (n = 50) were compared using Welch’s t-tests. The two groups did not differ in age (*t*_(98.3)_ = 1.07, *P* = 0.29), age of onset of musical training (*t*_(98.9)_ = −0.84, *P* = 0.40), cumulative musical training (*t*_(95.19)_ = 0.97, *P* = 0.33), crystallized intelligence (*t*_(96.4)_ = −1.48, *P* = 0.14), and fluid intelligence (*t*_(96.7)_ = −1.78, *P* = 0.08). As predicted, AP musicians had a substantially higher tone-naming score than RP musicians (*t*_(99)_ = 13.53, *P* < 10^−15^). There was a trend towards a higher musical aptitude in AP musicians as quantified by the AMMA total score (*t*_(97.2)_ = 1.99, *P* = 0.05). Follow-up analyses of the AMMA subscores showed that this difference was driven by a slightly higher tonal score in AP musicians (*t*_(96.5)_ = 2.27, *P* = 0.03), but there was no difference regarding the rhythm score (*t*_(98.0)_ = 1.42, *P* = 0.16). Descriptive statistics of participant characteristics are given in Table 1.

The in-scanner behavioral measures were analyzed using a mixed-design ANOVA with a within-subject factor Condition (Listening vs. Labeling) and a between-subject factor Group (AP vs. RP). As shown in Figure 2A, the mixed-design ANOVA of the response accuracy revealed an interaction between the factors Group and Condition (*F*_(1,99)_ = 8.37, *P* = 0.005, η^2^_G_ = 0.02). The difference in response accuracy between the two conditions (Labeling minus Listening) was smaller in AP than in RP musicians (Welch’s t-test, *t*_(79.1)_ = 2.88, *P* = 0.005, *d* = 0.57). Furthermore, this difference correlated with the tone-naming score (*r* = 0.41, *P* < 0.001). On average, the response accuracy was higher in the Listening condition than in the Labeling condition, so this correlation indicates a smaller difference for participants with a higher tone-naming score (see Figure 2C). Additional follow-up analyses showed a higher response accuracy for AP musicians in the Labeling condition (Welch’s t-test, *t*_(73.4)_ = 2.88, *P* = 0.005, *d* = 0.57), but not in the Listening condition (Welch’s t-test, *t*_(87.7)_ = 1.10, *P* = 0.28, *d* = 0.22). As shown in Figure 2B, the mixed-design ANOVA of the response time revealed a Group x Condition interaction (*F*_(1,99)_ = 8.85, *P* = 0.004, η^2^_G_ = 0.01). The condition difference in response time was smaller in AP musicians (Welch’s t-test, *t*_(95.6)_ = −2.97, *P* = 0.004, *d* = 0.59). Again, this difference correlated with the tone-naming score (*r* = −0.31, *P* = 0.002) (see Figure 2D). Descriptive statistics of the in-scanner behavioral measures are given in Table 2.

**Figure 2.**
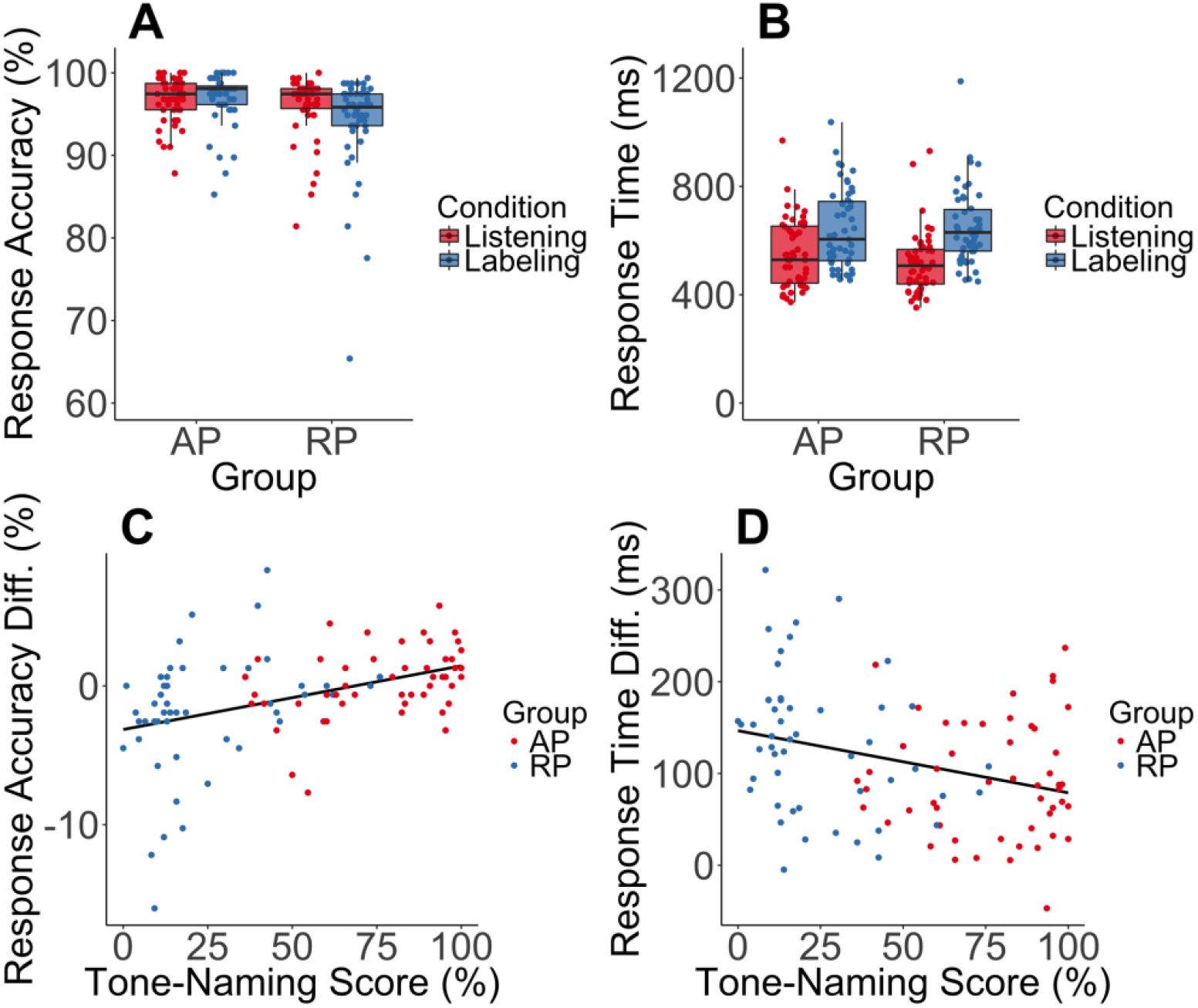
In-scanner behavioral measures (response accuracy and response time). (*A*) Interaction between Group (AP vs. RP) and Condition (Listening vs. Labeling) as revealed by a mixed-design ANOVA of the response accuracy (*F*_(1,99)_ = 8.37, *P* = 0.005, η^2^_G_ = 0.02). The interaction is characterized by smaller differences between Listening and Labeling in AP musicians than RP musicians (*t*_(79.1)_ = 2.88, *P* = 0.005, *d* = 0.57). Additionally, AP musicians demonstrated higher response accuracy in the Labeling condition (*t*_(73.4)_ = 2.88, *P* = 0.005, *d* = 0.57). (*B*) Group x Condition interaction as revealed by a mixed-design ANOVA of response time (*F*_(1,99)_ = 8.85, *P* = 0.004, η^2^_G_ = 0.01), again characterized by smaller condition differences in AP musicians (*t*_(95.6)_ = −2.97, *P* = 0.004, *d* = 0.59). (*C*) Correlation between the condition difference in response accuracy (Labeling minus Listening) and tone-naming score (*r* = 0.41, *P* < 0.001). Note that the positive correlation indicates a smaller difference for participants with a higher tone-naming score. (*D*) Correlation between the condition difference in response time and tone-naming score (*r* = −0.31, *P* = 0.002). AP = absolute pitch, RP = relative pitch.

**Table 2.** In-scanner behavioral measures. Measures given as mean ± standard deviation.

|  | AP musicians | RP musicians |
| --- | --- | --- |
| Response accuracy Listening | 96.69 ± 2.68 % | 95.97 ± 3.82 % |
| Response accuracy Labeling | 96.88 ± 3.14 % | 94.12 ± 6.04 % |
| Response time Listening | 550.23 ± 125.60 ms | 516.20 ± 114.61 ms |
| Response time Labeling | 642.33 ± 145.40 ms | 649.29 ± 139.13 ms |

Abbreviations: AP = absolute pitch, RP = relative pitch

### BOLD Signal Changes

The BOLD signal changes were analyzed using a voxel-wise GLM in combination with a second-level mixed factorial design., Using the mask restricting the search space to brain regions previously associated with AP or RP, we found a Group x Condition interaction which was characterized by smaller BOLD signal condition differences in AP musicians, paralleling the in-scanner behavioral measures. As shown in Figure 3A, this interaction was detected in three frontal clusters (see Table 3 for details). FWE-corrected *P* values (*P_FWE_*) and the number of voxels (*k*) of clusters are given in brackets. The clusters were localized in the right IFG, pars opercularis (*P_FWE_* < 0.001, *k* = 407) and the left IFG, pars opercularis (*P_FWE_* = 0.003, *k* = 169). A third cluster was localized in the presupplementary motor area (preSMA) of the dorsomedial prefrontal cortex (*P_FWE_* = 0.005, *k* = 141). The exploratory whole-brain analysis for the Group x Condition interaction yielded virtually identical clusters with the same maxima. These clusters were slightly more extended than in the restricted analysis. For full transparency, we made the unthresholded *t*-maps of the whole-brain analyses available on NeuroVault (Gorgolewski et al. 2015), https://neurovault.org/collections/4906/.

**Figure 3.**
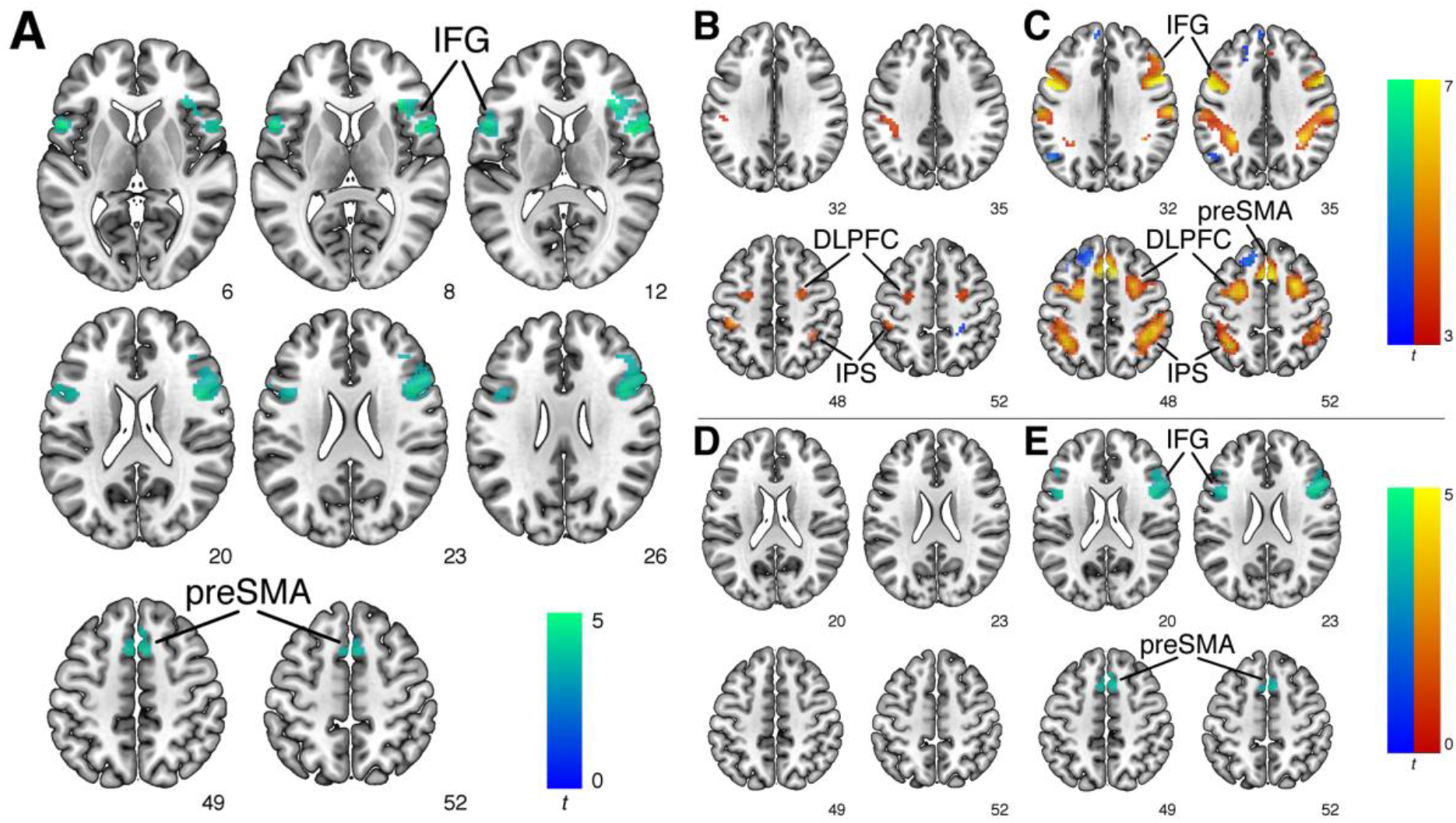
Results of the GLM analysis. (*A*) Group x Condition interaction characterized by smaller condition differences in AP musicians in the right inferior frontal gyrus (IFG), pars opercularis (*P_FWE_* < 0.001, *k* = 407), left IFG, pars opercularis (*P_FWE_* = 0.003, *k* = 169), and presupplementary motor area (preSMA) of the dorsomedial prefrontal cortex (*P_FWE_* = 0.005, *k* = 141). Cold colors indicate AP < RP. (*B*) Follow-up analysis within AP musicians revealed increases during Labeling in bilateral intraparietal sulcus (IPS) and bilateral dorsolateral prefrontal cortex (DLPFC). Hot colors indicate Labeling > Listening and cold colors indicate Listening > Labeling. (*C*) Follow-up analysis within RP musicians revealed similar increases during Labeling in bilateral IPS and bilateral DLPFC and unique increases in the bilateral IFG and the preSMA. Hot colors indicate Labeling > Listening and cold colors indicate Listening > Labeling. (*D*) Follow-up analysis within the Listening condition revealed no group differences. (*E*) Follow-up analysis within the Labeling condition revealed equivalent clusters to the Group x Condition interaction in the right IFG (*P_FWE_* < 0.001, *k* = 312), the left IFG (*P_FWE_* = 0.003, *k* = 195), and the preSMA (*P_FWE_* = 0.005, *k* = 134). Cold colors indicate AP < RP. AP = absolute pitch, RP = relative pitch.

**Table 3.**
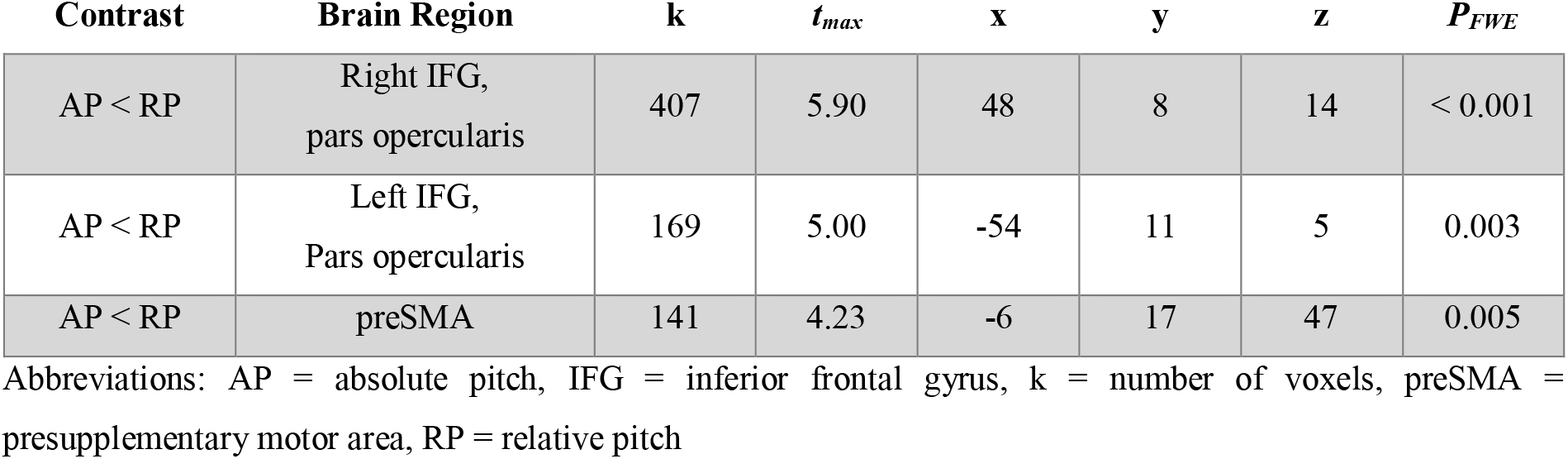
Group x Condition interaction of BOLD signal. The coordinates (x, y, z) are in MNI space. The clusters are ordered according to their size.

| Contrast | Brain Region | k | $t_{max}$ | x | y | z | $P_{FWE}$ |
| --- | --- | --- | --- | --- | --- | --- | --- |
| AP < RP | Right IFG,<br>pars opercularis | 407 | 5.90 | 48 | 8 | 14 | < 0.001 |
| AP < RP | Left IFG,<br>Pars opercularis | 169 | 5.00 | -54 | 11 | 5 | 0.003 |
| AP < RP | preSMA | 141 | 4.23 | -6 | 17 | 47 | 0.005 |

Abbreviations: AP = absolute pitch, IFG = inferior frontal gyrus, k = number of voxels, preSMA = presupplementary motor area, RP = relative pitch

As shown in Figure 3B and 3C, using the restricted search space, follow-up analyses within each group separately revealed similar BOLD signal differences between the two conditions with the exception of the three clusters described above (bilateral IFG, preSMA). In the bilateral IFG and the preSMA, only RP musicians showed increased BOLD signal in the Labeling condition. In addition, both groups showed increases in the bilateral intraparietal sulcus (IPS) and the bilateral DLPFC (see Table 4). These increases were stronger and more distributed in RP musicians, again indicating larger condition differences. Further follow-up analyses within each condition revealed that there were no group differences in the Listening condition (see Figure 3D). In contrast, AP musicians showed lower BOLD signal in the Labeling condition in the right IFG (*P_FWE_* < 0.001, *k* = 312), the left IFG (*P_FWE_* = 0.003, *k* = 195), and the preSMA (*P_FWE_* = 0.005, *k* = 134). These clusters were equivalent to the clusters of the Group x Condition interaction (see Figure 3E and Table 5). The whole-brain analysis yielded again virtually identical results with slightly extended clusters. Unthresholded t-maps of the whole-brain follow-up analyses are available on NeuroVault (https://neurovault.org/collections/4906/).

**Table 4.** Condition differences in BOLD signal within each group. The coordinates (x, y, z) are in MNI space. The clusters are ordered according to the contrast, the group, and the cluster size.

| Group | Contrast | Brain Region | k | $t_{max}$ | x | y | z | $P_{FWE}$ |
| --- | --- | --- | --- | --- | --- | --- | --- | --- |
| AP | Labeling > Listening | Left IPS | 101 | 5.37 | -42 | -31 | 50 | 0.009 |
| AP | Labeling > Listening | Left DLPFC | 69 | 4.56 | -27 | -7 | 47 | 0.01 |
| AP | Labeling > Listening | Right DLPFC | 41 | 4.59 | 27 | -4 | 50 | 0.03 |
| AP | Labeling > Listening | Right IPS | 29 | 4.51 | 36 | -43 | 44 | 0.04 |
| RP | Labeling > Listening | Left IFG, pars opercularis, preSMA, Left DLPFC | 1290 | 9.85 | -57 | 8 | 23 | < 0.001 |
| RP | Labeling > Listening | Right IFG, pars opercularis, Right DLPFC | 1053 | 10.95 | 54 | 8 | 14 | < 0.001 |
| RP | Labeling > Listening | Left IPS | 622 | 7.01 | -33 | -46 | 38 | < 0.001 |
| RP | Labeling > Listening | Right IPS | 595 | 7.14 | 39 | -46 | 44 | < 0.001 |
| AP | Listening > Labeling | Right SPL | 58 | 4.48 | 30 | -34 | 62 | 0.02 |
| RP | Listening > Labeling | Left DLPFC | 156 | 5.05 | -21 | 38 | 41 | 0.004 |
| RP | Listening > Labeling | Left frontal pole | 91 | 5.93 | -9 | 56 | 23 | 0.01 |
| RP | Listening > Labeling | Right SPL | 58 | 4.72 | 12 | -46 | 71 | 0.02 |
| RP | Listening > Labeling | Left angular gyrus | 45 | 4.97 | -51 | -64 | 32 | 0.03 |

Abbreviations: AP = absolute pitch, DLPFC = dorsolateral prefrontal cortex, IPS = intraparietal sulcus, IFG = inferior frontal gyrus, k = number of voxels, preSMA = presupplementary motor area, RP = relative pitch, SPL = superior parietal lobule.

**Table 5.** Group differences in BOLD signal in the Labeling condition. The coordinates (x, y, z) are in MNI space. The clusters are ordered according to their size.

| Contrast | Brain Region | k | $t_{max}$ | x | y | z | $P_{FWE}$ |
| --- | --- | --- | --- | --- | --- | --- | --- |
| AP < RP | Right IFG,<br>pars opercularis | 312 | 4.63 | 45 | 11 | 23 | < 0.001 |
| AP < RP | Left IFG,<br>Pars opercularis | 195 | 4.17 | -42 | 8 | 23 | 0.003 |
| AP < RP | preSMA | 134 | 4.69 | 9 | 23 | 44 | 0.005 |

Abbreviations: AP = absolute pitch, IFG = inferior frontal gyrus, k = number of voxels, preSMA = presupplementary motor area, RP = relative pitch

### Group Decoding by Searchlight Analysis

In addition to the voxel-wise GLM, we used searchlight analysis to localize BOLD signal patterns which differentiate between the two groups (Kriegeskorte et al. 2006). For the main analysis, we used the difference in BOLD signal patterns between the two conditions as the input. As shown in Figure 4A, group status could be decoded in the left IFG, pars triangularis (*P_FWE_* = 0.01, *k* = 29). The mean classification accuracy within the cluster was 72.5%. In comparison to the left IFG cluster from the GLM Group x Condition interaction, this cluster was located more anteriorly on the IFG. Follow-up analyses were performed with the patterns of each condition separately. Analogous to the GLM analysis, group status could not be decoded based on patterns in the Listening condition. In contrast, group status could be decoded based on Labeling patterns in the preSMA (*P_FWE_* < 0.001, *k* = 81, mean classification accuracy = 70.6%). This cluster substantially overlapped with the preSMA cluster from the GLM (see Figure 4A). However, a complete overlap should not be expected, because searchlight analysis is known to cause slight distortions in the localization (Etzel et al. 2013).

**Figure 4.**
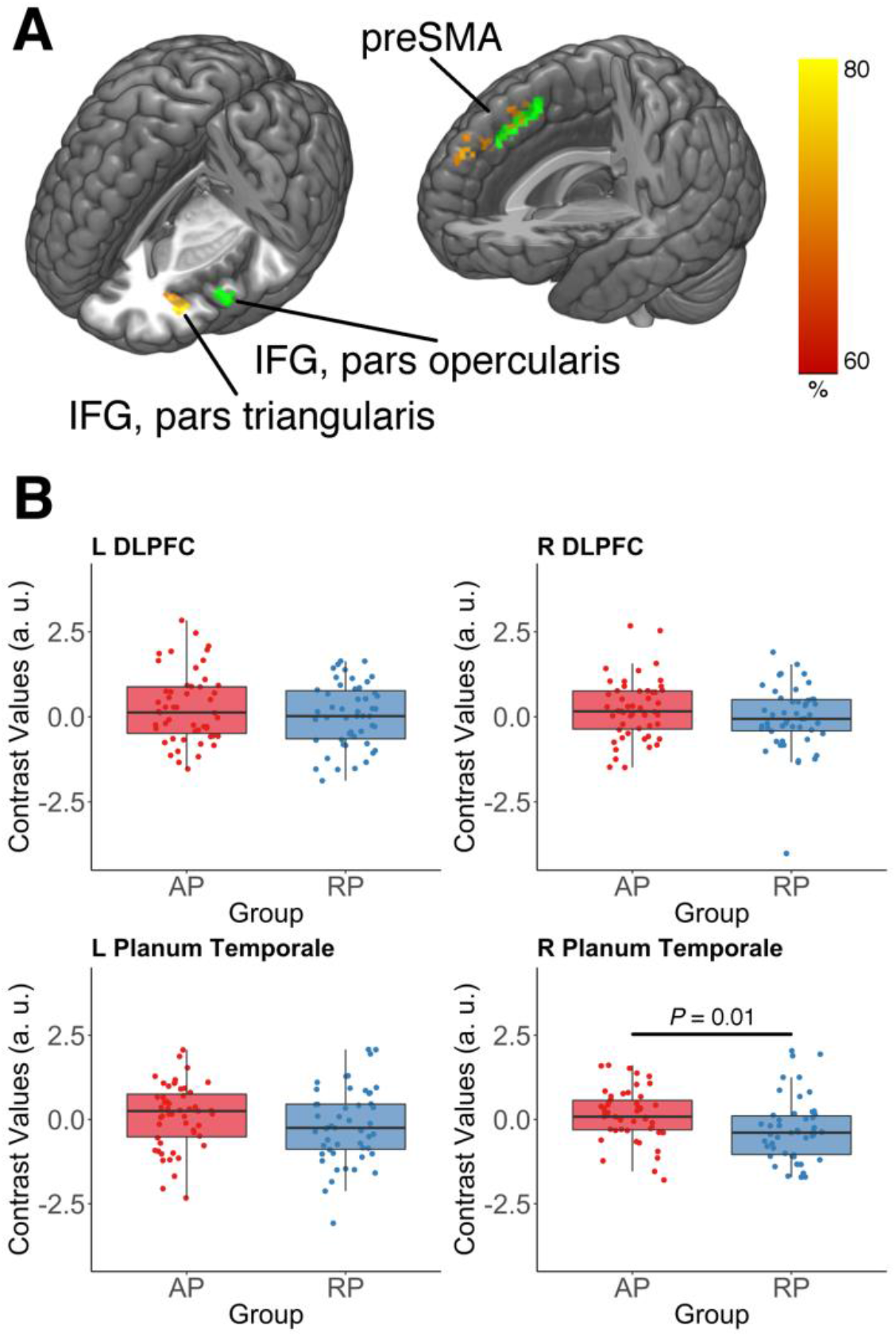
Results of the searchlight analysis and the ROI analysis. (*A*) Left: Group status could be decoded in the left inferior frontal gyrus (IFG), pars triangularis (*P_FWE_* = 0.01, *k* = 29) based on the difference in BOLD signal patterns between Listening and Labeling (shown in red-yellow). The cluster is located more anteriorly on the IFG compared to the Group x Condition cluster from the GLM analysis (shown in green). Right: Group status decoding in the presupplementary motor area (preSMA, *P_FWE_* < 0.001, *k* = 81) based on patterns in the Labeling condition (shown in red-yellow). There is substantial overlap with the preSMA cluster revealed by the GLM group comparison during Labeling (shown in green). Hot colors represent the classification accuracy. (*B*) AP musicians show higher mean BOLD signal chances during Listening in the right planum temporale (*t*_(94.6)_ = 2.66, *P* = 0.01, *d* = 0.53), but not in the left planum temporale or the bilateral dorsolateral prefrontal cortex (DLPFC). AP = absolute pitch, RP = relative pitch.

### Regional Mean BOLD Signal Changes

Finally, we extracted the mean BOLD signal changes from a priori defined ROIs. The bilateral planum temporale and the bilateral DLPFC were used as ROIs as these regions have previously been associated with AP processing (Schlaug et al. 1995; Zatorre et al. 1998; Keenan et al. 2001; Ohnishi et al. 2001; Bermudez and Zatorre 2005; Wilson et al. 2009; Wengenroth et al. 2014). It has been proposed that AP musicians automatically retrieve the pitch-label association from long-term memory when confronted with tones (Itoh et al. 2005). Therefore, the group comparison of mean BOLD signal changes was only performed in the Listening condition (Zatorre et al. 1998; Ohnishi et al. 2001; Itoh et al. 2005). As described above, we did not find group differences during Listening with the voxel-wise GLM analysis and the searchlight analysis. However, these analyses may miss subtle effects related to the automatic retrieval because of their conservative correction for multiple comparisons (Poldrack 2007). As shown in Figure 4B, AP musicians showed increased mean BOLD signal in the right planum temporale (Welch’s t-test, *t*_(94.6)_ = 2.66, *P* = 0.01, *d* = 0.53), but not in the left planum temporale, the left DLPFC, and the right DLPFC (all *P* > 0.10). The exploratory ROI analysis of bilateral Heschl’s gyrus did not reveal group differences in mean BOLD signal in the left Heschl’s gyrus (*P* = 0.20). Also, the mean BOLD signal in the right Heschl’s gyrus did not significantly differ between the groups (*P* = 0.09), although there was descriptively a tendency towards higher BOLD signal in AP musicians associated with a small effect size (*d* = 0.34).

## Discussion

In this study, we investigated AP and RP processing in the human brain using task-based fMRI in a large sample of musicians. The GLM analysis revealed smaller BOLD signal differences between Listening and Labeling in AP musicians than in RP musicians. The smaller differences between the conditions were driven by lower BOLD signals in AP musicians during Labeling in the left- and right-sided pars opercularis of the IFG and the preSMA. The in-scanner behavioral measures (response accuracy and response time) mirrored the fMRI data by showing smaller differences between Listening and Labeling in AP musicians. Using MVPA, we found that group status could be decoded in the left-sided pars triangularis of the IFG based on the difference in BOLD signal patterns between Listening and Labeling. Furthermore, group decoding was also possible in the preSMA based on BOLD signal patterns obtained in the Labeling condition. Lastly, the ROI analysis revealed a higher mean BOLD signal in AP musicians during Listening in the right planum temporale which was not detected by the GLM analysis and the MVPA.

The IFG is an important target region for auditory information which is propagated from the auditory cortex to the IFG along the ventral stream (the “what” pathway) of auditory processing (Rauschecker and Scott 2009). In this context, the IFG has been repeatedly linked with auditory working memory functions (Schulze et al. 2018). More specifically, the IFG has been associated with working memory for pitch, as shown by both PET and fMRI studies (Zatorre et al. 1994; Gaab et al. 2003). In this study, we observed BOLD signal increases in RP musicians bilaterally in the IFG during Labeling. This increase was not observable in AP musicians. As RP musicians need to use their RP ability to successfully complete the task, it is plausible that the signal increase in the IFG reflects pitch working memory processes as an important aspect of RP processing (McDermott and Oxenham 2008). This interpretation is fully in line with the results of the PET study described in the introduction (Zatorre et al. 1998). In this study, RP musicians, but not AP musicians, showed CBF increases in IFG while they were labeling musical intervals. More evidence for the association between RP processing and working memory comes from a number of electrophysiological studies investigating the P300 component of the auditory event-related potential. The P300 presumably reflects the updating of auditory information in working memory. Several studies found an absent or reduced P300 component in AP musicians not relying on RP processing. In contrast, RP musicians show a normal P300 amplitude (Klein et al. 1984; Itoh et al. 2005).

Apart from being implicated in working memory, the IFG has been strongly associated with language functions. In the left hemisphere, the pars opercularis (Brodmann area 44) and the pars triangularis (Brodmann area 45) of the IFG are known as Broca’s area, a brain region traditionally associated with speech production, but also heavily involved in speech perception (Friederici 2011). In the right hemisphere, the IFG is linked to the perception of prosody (pitch changes in speech) (Buchanan et al. 2000). Therefore, the BOLD signal increases in RP musicians in bilateral IFG might reflect language-related processes. More concretely, the RP musicians might have engaged in covert articulation of the tone labels as a part of their strategy to label the tones. In contrast, it seems that the AP musicians do not rely on a verbal code to successfully complete the task. This is in accordance with behavioral evidence demonstrating non-verbal coding strategies in AP musicians (Zatorre and Beckett 1989), and fMRI evidence showing atypically similar BOLD signal in AP musicians during the perception of tonal and verbal stimulus material (Schulze et al. 2013).

Mirroring the bilateral IFG BOLD signal increases, the preSMA showed signal increases in RP musicians during Labeling. In addition, the BOLD signal patterns during Labeling in the preSMA contained information about group status. Thus, AP and RP processing were accompanied by differential BOLD signal patterns. The preSMA is anatomically connected to the IFG via the frontal aslant tract and has been implicated in speech production and processing (Catani et al. 2013). More importantly, the preSMA plays a key role in the auditory imagery of pitch (Lima et al. 2016). Auditory imagery generally refers to the generation of auditory information in the absence of sound perception. However, auditory imagery can also involve auditory information that is generated in addition to the currently perceived information. Consequently, RP musicians might have imagined the pitches of previously heard tones to determine the pitch of the current tone. This interpretation is in line with the anecdotal observation that RP musicians often covertly sing pitches in order to identify the musical intervals. It is important to note that the working memory and the language explanations of the IFG and preSMA involvement during Labeling are not mutually exclusive. There is evidence that largely overlapping brain regions are involved in auditory working memory for verbal material and non-verbal material, for example, pitches (Koelsch et al. 2009).

The results from the GLM analysis and the MVPA did not fully converge with regard to the localization of the group differences. Most notably, using MVPA, we found that group status could be decoded from BOLD signal patterns in the left-sided pars triangularis of the IFG whereas the GLM revealed BOLD signal differences in the pars opercularis. As mentioned above, these two regions constitute Broca’s area. In a previous study using MVPA, it was shown that BOLD signal patterns in Broca’s area contain speech-related information which was not detectable with GLM analysis (Lee et al. 2012). MVPA is more sensitive to information in fine-grained patterns which are preserved in unsmoothed fMRI data (Kriegeskorte and Bandettini 2007). At the same time, there has been a debate about whether or not Broca’s area should be divided into subareas executing different functions (Friederici 2011). Consequently, we propose that the BOLD signal patterns in the pars triangularis represent information about AP and RP on a smaller spatial scale. In contrast, the differences in the pars opercularis might be more homogeneous and therefore detectable by the GLM analysis. Further studies should elucidate the potentially differential roles of these two brain regions in pitch processing.

Although showing lower BOLD signal in the IFG and preSMA during Labeling, the AP musicians identified the tones more accurately than RP musicians. Therefore, AP processing might be more efficient than RP processing with regard to the use of neural resources. Neural efficiency has been discussed in relation to intelligence, where it has been proposed that more intelligent individuals show lower BOLD signal while performing cognitive tasks (Neubauer and Fink 2009). In this study, there were no group differences in psychometrically evaluated intelligence. Neural efficiency is often observed in tasks of low or moderate difficulty and predominantly in brain regions of the frontal cortex (Neubauer and Fink 2009). Both of these prerequisites are present in this study. The efficiency of AP processing might be related to the automatic retrieval of the pitch-label association which presumably occurs immediately after the pitch is encoded (Itoh et al. 2005). This process is often described as effortless (Deutsch 2013). RP requires more processing steps because after the encoding, the pitch needs to be compared to a previous pitch held in working memory and subsequently, the exact interval between those two pitches needs to be determined. One might speculate that the presumed neural efficiency of AP processing could be a reason for its continued existence throughout human evolution despite its negligible role in music and speech perception (McDermott and Oxenham 2008). On the other hand, it could also be argued that AP musicians did not use the IFG and preSMA at all during Labeling, and thus, the notion of more efficient neural processing might be misplaced, as AP musicians might have used different brain regions than RP musicians and not the same regions more efficiently (see Neubauer and Fink 2009). Following this line of reasoning, AP musicians may have relied on different cognitive processes during Labeling than RP musicians. However, there are two lines of evidence that speak against the AP-specific use of fundamentally different neural resources in the Labeling condition: First, from the unthresholded statistical map displaying differences in BOLD signal between Listening and Labeling within AP musicians, one can observe that AP musicians did, to some extent, activate the bilateral IFG and the preSMA more during Labeling than during Listening (see https://neurovault.org/images/117517/). Thus, they actually used the same, or at least similar, brain regions as RP musicians during Labeling. Second, recent behavioral studies have demonstrated that AP processing and higher cognitive functions (e.g., working memory) are more closely related than previously thought (Van Hedger et al. 2015; Van Hedger and Nusbaum 2018). Hence, it is possible that AP processing is not completely independent of higher cognitive functions but relies less on them than RP processing.

During Listening, the AP musicians showed larger BOLD signals than RP musicians in the right planum temporale. We observed this increase exclusively with the ROI analysis, so the effect seems to be spatially restricted and too subtle to be detected by analyses employing a conservative correction for multiple comparisons. As described in the introduction, the planum temporale has been associated with AP processing from the very beginning of neuroscientific AP research (Schlaug et al. 1995). It is part of the non-primary auditory cortex and has an important role in the processing of a diverse range of sounds (Griffiths and Warren 2002). In this study, the increase in signal was restricted to the right hemisphere. This finding is consistent with previous studies reporting anatomical differences in AP musicians in the right planum temporale (Keenan et al. 2001; Wilson et al. 2009; Wengenroth et al. 2014) and with an influential theory on the importance of the right hemispheric auditory cortex in music processing (Zatorre et al. 2002). However, its exact role in AP processing is still unclear. With regard to auditory processing in general, it has been proposed that the planum temporale matches incoming auditory information with information that is stored in templates which are not located in the planum temporale itself (Griffiths and Warren 2002). According to the two-component model, AP musicians possess long-term representations of pitches associated with meaningful labels. These representations could well be characterized as internal templates to which incoming information is matched (Levitin 1994; Levitin and Rogers 2005). Therefore, we propose that in AP musicians, incoming auditory information, more precisely the extracted pitch information, is matched with these internal pitch templates by computations performed in the right planum temporale. The templates themselves could be represented in more anterior regions of the right temporal lobe which are implicated in semantic memory (Binder and Desai 2011). In line with this idea, two recent studies investigating AP musicians have found evidence for differential structural and functional connectivity along the right-hemispheric ventral stream of auditory processing, especially in the planum polare which is located immediately anterior to Heschl’s gyrus (Kim and Knösche 2016, 2017). Thus, it will be interesting for future studies trying to consolidate the findings of AP-specific alterations in posterior and anterior secondary auditory cortices.

In contrast to the previously described PET study, we did not find group differences in the posterior DLPFC during Listening. In the PET study, the involvement of the DLPFC was attributed to the automatic retrieval of the pitch-label association in AP musicians (Zatorre et al. 1998). The current results do not support this interpretation. In both groups, we observed bilateral DLPFC BOLD signal increases during Labeling. These increases were accompanied by higher BOLD signal in the bilateral IPS, again in both groups. Both the DLPFC and the IPS are important parts of a network strongly linked to top-down attentional control (Corbetta and Shulman 2002). Therefore, it is possible that the DLPFC involvement is related to unspecific attentional processes rather than the specific retrieval of the pitch-label association.

In conclusion, the current results indicate a possible involvement of working memory, language-related processes, and auditory imagery in RP processing, mediated by the bilateral IFG and the preSMA. AP musicians do not show BOLD signal increases in the IFG and the preSMA during Labeling. At the same time, AP musicians label the tones with a higher accuracy. This suggests that AP might be an example of neural efficiency, which is characterized by higher behavioral performance in combination with a lower use of neural resources. Using MVPA, we detected differential BOLD signal patterns in the IFG and the preSMA. Therefore, these regions might contain information differentiating AP from RP on a small spatial scale. Finally, during Listening, the AP musicians show a specific signal increase in the right planum temporale, possibly reflecting the matching of pitch information with internal templates. Taken together, AP and RP musicians show diverging frontal activations during Labeling and, more subtly, differences in right auditory activation during Listening. The results of this study do not support the previously reported importance of the posterior DLPFC in associating a pitch with its label.

## Compliance with Ethical Standards

### Ethical approval

All procedures performed in studies involving human participants were in accordance with the ethical standards of the institutional and/or national research committee and with the 1964 Helsinki declaration and its later amendments or comparable ethical standards.

### Informed consent

Informed consent was obtained from all individual participants included in the study.

### Funding

This work was supported by the Swiss National Science Foundation (SNSF), grant no. 320030_163149 to LJ.

## Acknowledgements

This work was supported by the Swiss National Science Foundation (SNSF), grant no. 320030_163149 to LJ. We thank our research interns Anna Speckert, Chantal Oderbolz, Désirée Yamada, Fabian Demuth, Florence Bernays, Joёlle Albrecht, Kathrin Baur, Laura Keller, Marilena Wilding, Melek Haçan, Nicole Hedinger, Pascal Misala, Petra Meier, Sarah Appenzeller, Tenzin Dotschung, Valerie Hungerbühler, Vanessa Vallesi, and Vivienne Kunz for their invaluable help in data acquisition and research administration. Without their help, this research would not have been possible. Furthermore, we thank Anja Burkhard for her support within the larger absolute pitch project, Roger Luechinger and Jürgen Hänggi for their assistance in specifying the MRI sequences, Silvano Sele for helpful discussions regarding the searchlight analysis, and Carina Klein, Stefan Elmer, and all other members of the Auditory Research Group Zurich (ARGZ) for their valuable comments on the experimental procedure.

